# An evolutionary transcriptomics approach links CD36 to membrane remodeling in replicative senescence

**DOI:** 10.1101/294512

**Authors:** Marie Saitou, Darleny Y. Lizardo, Recep Ozgur Taskent, Alec Millner, Gunes Ekin Atilla-Gokcumen, Omer Gokcumen

## Abstract

Cellular senescence, the irreversible ceasing of cell division, has been associated with organismal aging, prevention of cancerogenesis, and developmental processes. As such, the evolutionary basis and biological features of cellular senescence remain a fascinating area of research. In this study, we conducted comparative RNAseq experiments to detect genes associated with replicative senescence in two different human cell lines and at different time points. We identified 841 and 900 genes (core senescence-associated genes) that are significantly up- and downregulated in senescent cells, respectively, in both cell lines. Our functional enrichment analysis showed that downregulated core genes are primarily involved in cell cycle processes while upregulated core gene enrichment indicated various lipid-related processes. We further demonstrated that downregulated genes are significantly more conserved than upregulated genes. Using both transcriptomics and genetic variation data, we identified one of the upregulated, lipid metabolism gene, *CD36* as an outlier. We found that overexpression of CD36 induces a senescence-like phenotype and, further, the media of CD36-overexpressing cells alone can induce a senescence-like phenotype in proliferating young cells. Moreover, we used a targeted lipidomics approach and showed that phosphatidylcholines accumulate during senescence in these cells, suggesting that upregulation of CD36 could contribute to membrane remodeling during senescence. Overall, these results contribute to the understanding of evolution and biology of cellular senescence and identify several targets and questions for future studies.

## Introduction

Cellular senescence is the irreversible arrest of cell division (Hayflick 1965). Most mammalian cells undergo senescence during the organismal lifespan. As such, cellular senescence is a fundamental feature of human cells. The internal and external factors stimulating cellular senescence are myriad and not completely understood (Campisi 2013). However, tumor suppressors p16 and/or p53 are ultimately activated and eventually result in permanent cell division arrest (Qian & Chen 2010).

The evolutionary implications of cellular senescence remain a fascinating area of study. It is clear from previous work that cell-cycle control pathways are central to the cessation of cell division (Ogryzko et al. 1996). However, the evolution of the triggers of these cascades are not resolved. This question is especially intriguing given that the total number of cell divisions, but not cell division rate, seem to vary drastically between species (Gillooly et al. 2012). The evolution of cellular senescence regulatory mechanisms are discussed within the context of antagonistic pleiotropy, where a beneficial genetic basis can be harmful in another aspect in organismal maintenance, which may result in senescence (Giaimo & d’Adda di Fagagna 2012), and disposable soma hypothesis, where a greater investment in development may result in diminished maintenance of organism, which leads to senescence (Kirkwood 2005).

This evolutionary framework has many implications for cancer formation and prevention and aging (Ohtani et al. 2009). For example, the senescence program may promote growth and tissue repair (Muñoz-Espín et al. 2013), while preventing harmful oncogenic activity. Moreover, senescent cells may eventually release pro-inflammatory and oncogenic factors, especially at later stages of organismal life (Campisi 2013). One explanation of these negative effects is the eventual inability of senescent cells to sequester the accumulating internal and external stresses (*e.g.*, oxidative stress) (Sanders et al. 2013). According to the disposable soma framework, this is because the detrimental effects of these stresses present themselves later in life, and may have lower fitness consequences.

Senescent cells secrete an array of biomolecules known as the senescence-associated secretory phenotype (SASP), with numerous and context-dependent effects on neighboring cells (Watanabe et al. 2017). As such, even though cellular senescence is a highly regulated cell-specific process, post-senescence processes have tissue and organismal level effects (Coppé et al. 2008). The exact content and functional implications of the SASP remain an active area of research. However, it is of note that senescent cells communicate via intercellular interactions (Biran et al. 2015), making this process self-perpetuating.

To elucidate the mechanisms by which replicative senescence is regulated and how senescent cells function after they stop dividing, we conducted an evolutionary transcriptomics analysis. This analysis led to the identification of *CD36*, which is involved in fatty acid uptake and metabolism (Jay et al. 2015; Xu et al. 2013), as a potential mediator of replicative senescence. Further investigations linked CD36 activity to SASP-mediated senescence in proliferating young cells.

## Results and Discussion

### Genes involved in replicative senescence are associated with organismal aging

In an earlier study, we characterized the genes that are altered in expression during replicative senescence in BJ cells (human foreskin fibroblast) (Lizardo et al. 2017). We found that the expressional changes in lipid-related genes were significantly higher as compared to the rest of the transcriptome, placing the regulation of lipids as a central pathway during replicative senescence. In addition, recent studies have shown cell-to-cell variability among senescent cells ((Wiley et al. 2017).

In this study, we aimed to refine transcriptome data to determine the “core” genes involved in replicative senescence by studying two different cell lines with different tissue origins. Specifically, we used BJ and MRC-5 (human lung fibroblast) cell lines, which are routinely used as models of replicative senescence (Shiozawa-West et al. 2015; Lizardo et al. 2017; Burrows et al. 2010). We cultured these cell lines from an early population doubling (PD~2 for BJs and PD~24 for MRC-5s) until they naturally reached their proliferative capacity (**Figure S1A)** and displayed an increased β-galactosidase activity (**Figure S1B**). To elucidate the changes in transcription as the cells senesced, we collected cells at three time points: early (PD=15 and 31 for BJ and MRC-5), mid-population (PD=31 and 46 for BJ and MRC-5), and old population (PD=42 and 67 for BJ and MRC-5). We conducted comparative RNAseq experiments to compare the changes in the transcriptome in a pairwise manner.

Unguided clustering analyses of the transcriptome without any *a priori* input resulted in distinct clustering of both cell-type and population-doubling stages based on the principal component analysis (**Figure 1A**) and hierarchical clustering (**Figure 1B**). Next, to detect genes that are involved in replicative senescence independent of the cell type, we compared the gene expression levels between young, mid-population, and old stages. Based on this analysis, we identified thousands of transcripts that show significant changes during replicative senescence in both cell lines (**Table S1, Figure 1C**). Using this dataset, we found that there were significantly more genes that showed expression changes in the same trend (*i.e.*, up or down-regulated) over population doubling in both cell lines (shared genes) as compared to random expectations from all the protein-coding genes (p<0.0001, odds ratio =1.75, Fisher’s Exact Test). This was concordant with our principal component and clustering analyses described above (**Figure 1A-B**) and indicate that these shared genes are likely involved in fundamental replicative senescence, rather than cell line-specific, processes. Based on this insight, we focused our analyses on these shared genes (core senescence-associated genes, **Figure 1C**) from this point on.

**Figure 1.**
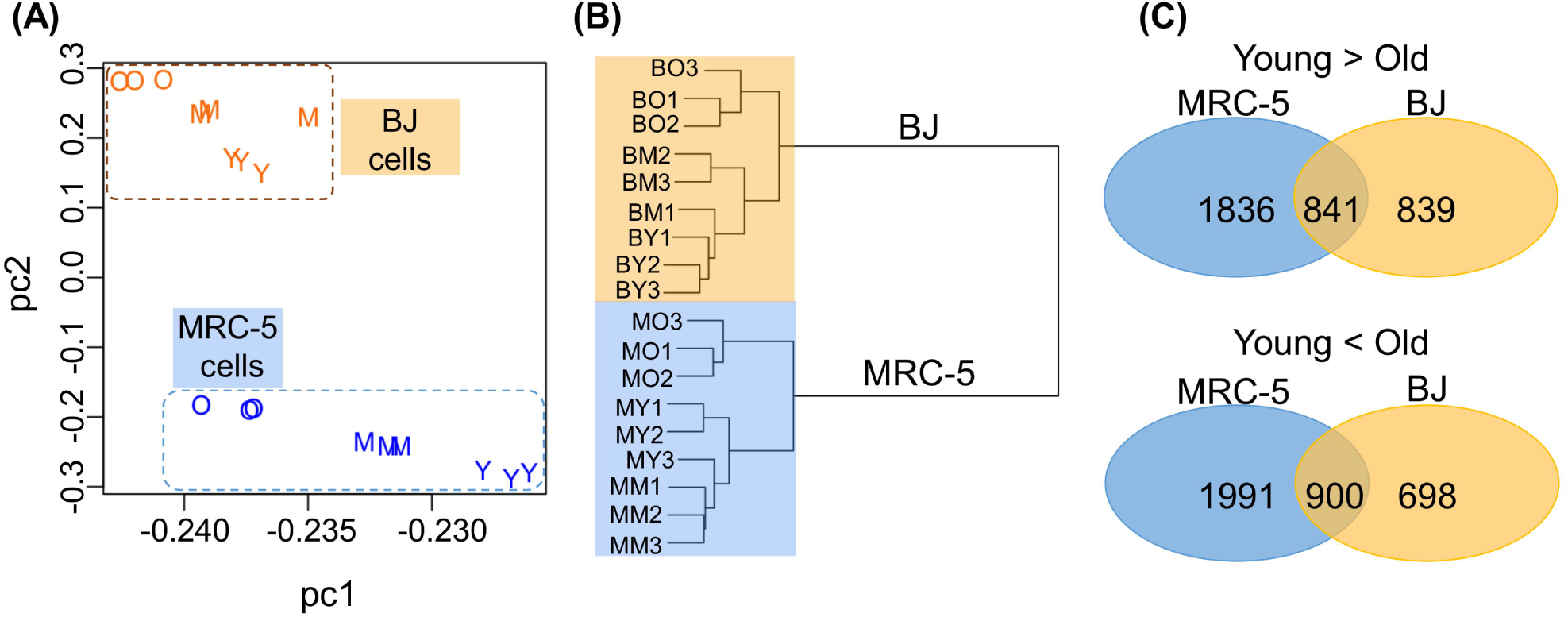
Characterization of replicative senescence by gene expressions. (A) Principal Component Analysis of all the genes of the different time points collected in MRC-5s and BJs. The first two principal components (PC1: 0.89 PC2: 0.069) are shown. Colors indicate cell lines (orange = BJs, blue = MRC-5s) and letters indicate age (“O” = old, “M” = mid-population, and “Y” = young). (B) Clustering dendrogram by transcriptomic data of the different time points collected in MRC-5s and BJs is shown. Spearman’s correlation of all the detected gene expression data of each cell was used. The dendrogram was drawn without any *a priori* clustering. Abbreviations: BY: young BJs, BM: mid-population BJs, BO: old BJs, MY: young MRC-5s, MM: mid-population MRC-5s, MO: old MRC-5s. (C) Venn diagram of genes that showed significant (*p*<0.0001, p-value adjusted for multiple testing using Benjamini-Hochberg method) fold change of expression between young and old in the two cell lines (BJ and MRC-5) and both cell lines.

Using this dataset, we tested if there were enriched genes reported with aging-associated traits to genes within the genome-wide association studies (GWAS) in core senescence-associated genes compared to the GWAS-negative protein-coding genes (**Table S2**). We observed an enrichment of GWAS-positive genes in genes that were significantly up- and down-regulated during replicative senescence (*p*-value = 1.16E-3, odds ratio = 1.64, Fisher’s Exact Test). These genes are prime candidates for further studies to determine the mechanistic links between replicative senescence and organismal aging.

### Signatures of a rapid shift towards senescence in cell populations

Our measurements of cellular senescence were at the cell population level as we assessed the changes in β-galactosidase activity in a given population of cells, as opposed to single-cell measurements. Since we studied three time points along the growth curve with detailed expression data, we asked whether the expression changes associated with replicative senescence gradually increased over time, or if there was a sudden population level, perhaps a “cliff-like”, change where changes rapidly occurred in the culture at a later stage. To test this, we categorized the longitudinal pattern of observed expression changes. This allowed us to discriminate between transcripts that increased (or decreased) over time, those that increased (or decreased) early (*i.e.*, from young to mid-population) or late (*i.e.*, from mid-population to old), and transcripts that exhibited a variable trend over time (*i.e.*, significantly increased early but decreased in later time points) (**Figure 2**).

**Figure 2.**
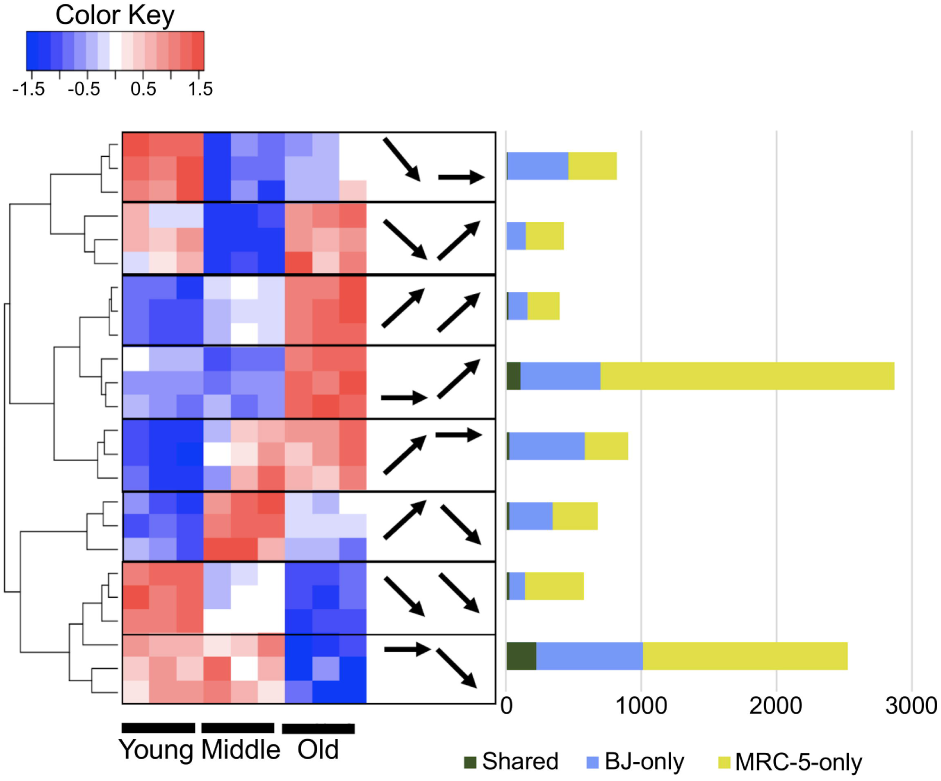
The expression change in three time points in the two cell lines. The heatmap shows relative gene expression of three representative genes (three rows) at each time point (red: higher expression, blue: lower expression). We categorized eight expression change patterns over three time points. Arrows indicate significant upregulation, downregulation or no significant change of gene expression between each time point. The bar graph represents the observed gene number in each of the eight expression pattern categories: genes observed in BJ cells (yellow), MRC-5 cells (blue), and both cell lines (dark green). Abbreviations: Young: Y, Mid-population: M, Old: O.

To visualize these categories, we plotted normalized gene expression for genes that showed significant differences between any two time points in either BJ or MRC-5 cell lines (**Figure 2**). The late stage changes in gene expression (from mid-population to old) explained majority of the observed transcriptomic changes during replicative senescence independent of the cell line. This result was also concordant with the decreased rate of population doubling that occurred in the late stages of replicative senescence.

### Genes involved in cell cycle processes were downregulated while genes involved in lipid metabolism were upregulated during replicative senescence

Next, to investigate the functional categories of genes associated with replicative senescence, we conducted Gene Ontology (GO) analysis (Ashburner et al. 2000; The Gene Ontology Consortium 2017) using Gorilla (Eden et al. 2009; Eden et al. 2007). We found that downregulated genes were primarily involved in cell cycle processes (FDR q-value=6.51E-28, **Figure 3**). Remarkably, 37 out of 841 downregulated genes in old cells were related to chromatin assembly. This observation is concordant with earlier studies where changes in chromatin organization have been implicated as a hallmark of senescence (Criscione et al. 2016). When we conducted a similar analysis to genes upregulated during replicative senescence, we found that the enrichment overwhelmingly indicated various lipid related processes, the top hit being “vesicle-mediated transport” (FDR q-value= 7.31E-6, **Figure 3, Table S3**). These findings fit well with previous studies, including ours (Lizardo et al. 2017), and other reports linking lipid regulation to senescence (Venable et al. 1995; Marthandan et al. 2016), suggesting important functional implications during this process. Moreover, similar functional categories were highlighted in organismal aging studies as well (Turan et al. 2018; Jobson et al. 2010).

**Figure 3.**
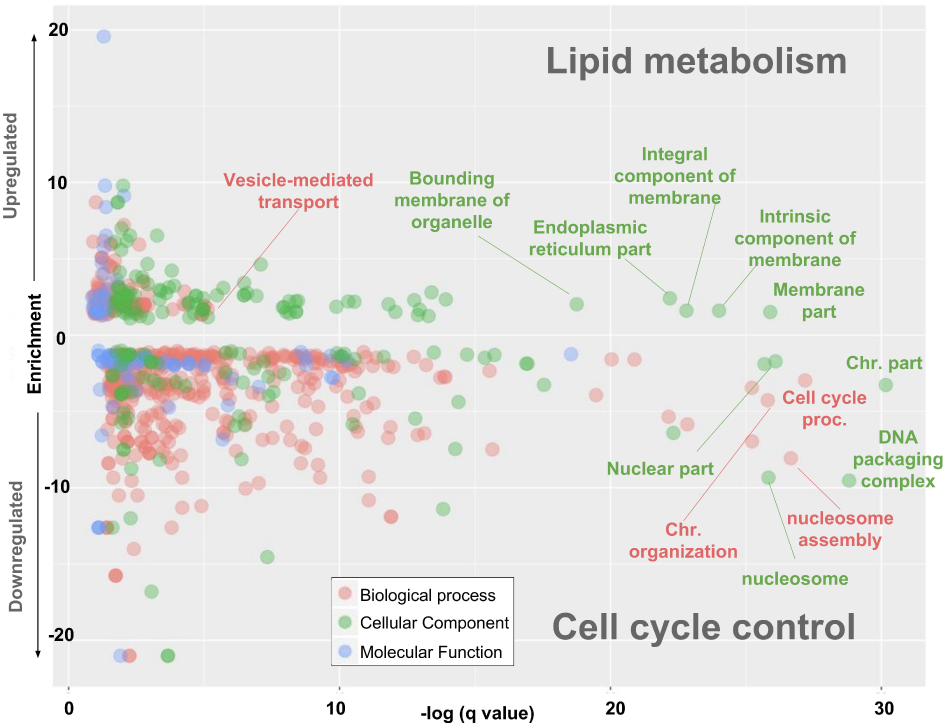
GO analysis of genes up-/down-regulated during replicative senescence. Each dot indicates an enriched GO category. X-axis shows -log q value (p-value corrected for multiple correction) of this enrichment. Y-axis shows the fold-change to all the protein-coding genes where positive values showed enrichments and negative values indicate depletion. The color indicates the three ontological categories (Biological Process, Cellular Component, Molecular Function).

### Evolutionary analysis of replicative senescence associated genes point to CD36 as a potential candidate for positive selection

As articulated by Campisi (Campisi 2013), the evolutionary forces that shape cellular senescence are complex. One previously articulated notion is that cellular senescence is a highly regulated cellular process akin to apoptosis (Coppé et al. 2008). Under this scenario, the expectation is that genes that trigger and maintain cellular senescence should be evolving under negative selection, which remove harmful mutations from a population during evolution. Based on the GO analysis of our transcriptomics data, we hypothesized, as we described above, that the downregulated genes might contribute to growth arrest. Thus, we expected these downregulated genes to be evolving under strong negative selection.

To test this, we measured the probability of being intolerant of homozygous loss-of-function variants (pLI value (Lek et al. 2016)) and impact of background selection (b-value (McVicker et al. 2009)) of all the protein-coding genes. The former is a measure comparing variation among healthy humans and the latter is a measure of negative selection based on datasets from multiple primate genomes. We then compared the level of negative selection between upregulated, downregulated, and all other protein-coding genes (**Figure 4A-B**). As expected, this analysis showed that up-/down-regulated genes are significantly less tolerant to loss of function variants (p<0.001, Mann-Whitney U test) and showed significantly higher signature of background selection, a measure of cross-species negative selection (p<0.001, Mann-Whitney U test), as compared to genes that are not involved in replicative senescence. This data further contributes to the notion that the cellular machinery that regulates replicative senescence is highly regulated and conserved. In other words, replicative senescence is a fundamental aspect of mammalian cellular life.

**Figure 4.**
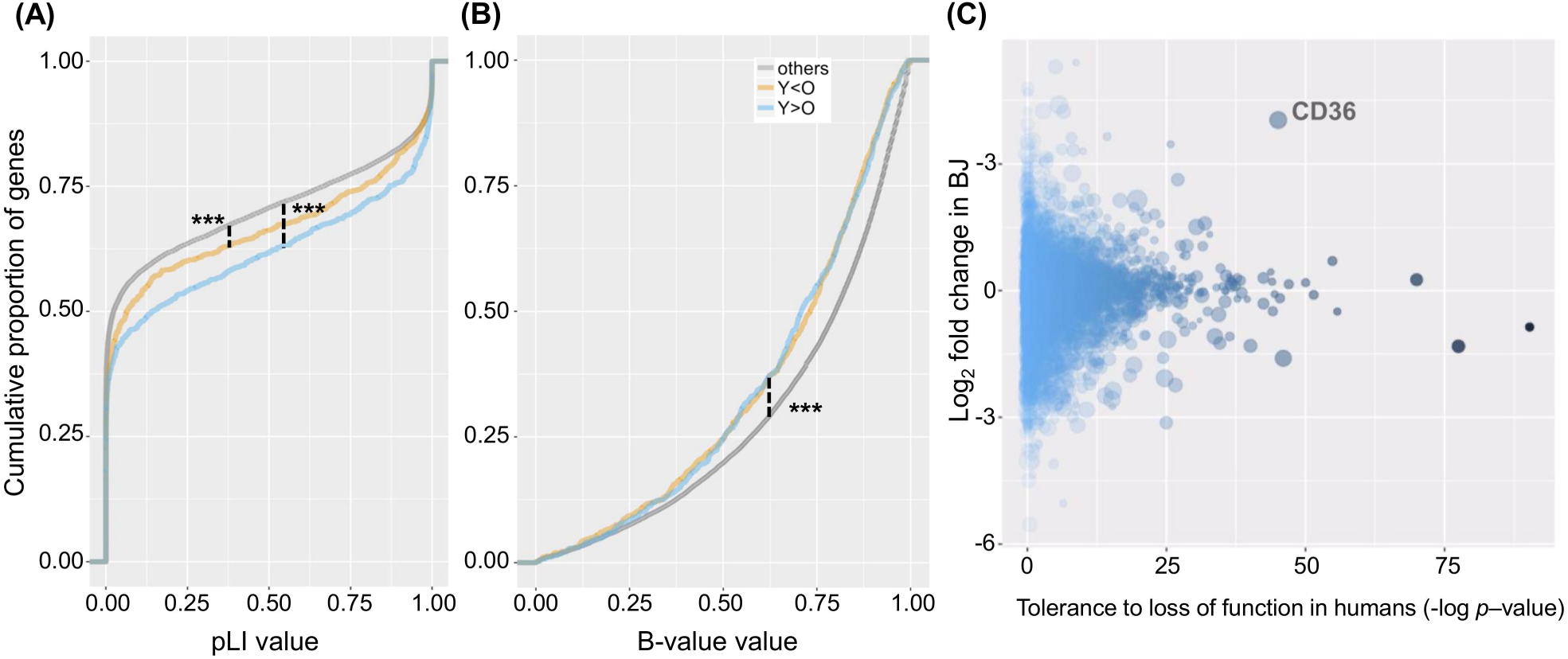
Characterization of genes associated with replicative senescence. (A) The cumulative fraction plot of pLI (the probability of being loss-of-function intolerant) value of genes upregulated in old cells (Y<O, orange), genes downregulated in old cells (Y>O, blue) and other protein-coding genes (gray). This measurement is obtained from ExAC database (Lek et al. 2016). (B) The cumulative fraction plot of b-value (the index of background selection between primate species) of genes in each category (McVicker et al. 2009). The color scheme is same as the previous panel. (C) X-axis shows the tolerance of genes in healthy human genomes (-log *p-*value of pLI), y-axis shows the log_2_ fold change between young and old in BJ cells and the size of the bubble indicates the log_2_ fold change between young and old in MRC-5 cells. The strength of the color shows the -log p-value of pLI. *CD36* showed upregulation in both old BJ and MRC-5 cells and tolerance to loss-of-function mutations in healthy human genomes.

Second, we considered the contemporary thinking about the evolution of cellular senescence (Campisi 2013) where some of the genes involved in replicative senescence may evolve in response to both intra- and extra-cellular stresses. Specifically, we surmised that some of these genes may have evolved under positive selection in adaptation to various stresses and show more tolerance to variation depending on environmental differences. These genes are also primary candidates to have antagonistic pleiotropic effects, a phenomenon extensively discussed within the context of cellular senescence (Bartholomew et al. 2009). Antagonistic pleiotropic effects explains that a beneficial genetic basis can be harmful in another aspect in organismal maintenance, which may result in senescence (Giaimo & d’Adda di Fagagna 2012). Based on our functional categorization, we argue that some of the upregulated, lipid-related genes involved in replicative senescence would be enriched in such variation tolerant, multi-functional genes. Indeed, we observed that upregulated genes in senescent cells show significantly more tolerance to mutations within humans as compared to downregulated genes (**Figure 4A**, p<0.05, Mann-Whitney U test).

Our previous work, along with others (Lizardo et al. 2017; Flor et al. 2017), showed that lipids play important roles in senescence. We therefore focused on genes related to lipid metabolism and found that *lipase member H (LIPH)*, a lipase that hydrolyzes phosphatidic acid, and *CD36 molecule (CD36)*, a scavenger receptor that has been shown to assist in fatty acid uptake, are the most upregulated genes during replicative senescence in both BJ and MRC-5 cell lines (**Table S1**). Further, a comparison of fold-expression changes during replicative senescence and tolerance to loss of function mutations revealed that *CD36* is a clear outlier (**Figure 4C**). It not only showed one of the highest upregulation in both BJ and MRC-5 cells during replicative senescence, but also had a higher number of functional mutations than expected within human populations. Indeed, previous population genetics work has identified that a loss of function variation of *CD36* is mostly observed in Central African populations and was increased in allele frequency likely through positive selection (Fry et al. 2009).

### Overexpression of CD36 resulted in senescence-like phenotype in young MRC-5 cells

CD36 is a scavenger receptor that has been implicated in various cellular processes, including long chain fatty acid binding and facilitating lipid transport into the cell as well as contributing to inflammatory responses (Silverstein & Febbraio 2009). CD36 activity has previously been linked to senescence (Yoon et al. 2016). We previously showed that during replicative senescence in BJ fibroblast cells there is a specific accumulation of polyunsaturated fatty acyl chain containing triacylglycerols as a result of CD36-mediated fatty acid uptake. There are different views on the involvement of CD36 in fatty acid metabolism, such as single/multiple fatty acid binding sites or enhancement of fatty acid metabolism (Jay et al. 2015; Xu et al. 2013), yet we still lack a clear understanding of the various roles this multifunctional protein fulfills.

In order to investigate CD36’s role during replicative senescence, we used transient transfection to overexpress FusionRed-CD36 in MRC-5 cells at population doubling 33. We observed a modest transfection efficiency in MRC-5 cells (~10%) (**Figure 5A, Figure S2**). We hypothesized that if the activation of CD36 is linked to senescence, its overexpression should result in a senescence-like phenotype. To test this, we transiently transfected MRC-5 cells with CD36 and tested for cellular senescence markers. We found that CD36 overexpression resulted in a significant and profound upregulation of β-galactosidase activity (**Figure 5B-C**) and slight increase in p16 expression (**Figure 5D**). This increase in β-galactosidase activity was unexpected given the modest transfection efficiency. That is, even though roughly 10% of the cells showed detectable overexpression of CD36 via confocal fluorescence microscopy imaging, almost all the cells in the population (90%) showed increased β-galactosidase activity (**Figure 5B-C**). These results indicate that CD36 upregulation not only mediates senescence in cells where it is overexpressed, but leads to initiation of senescence in the cell population as a whole.

**Figure 5.**
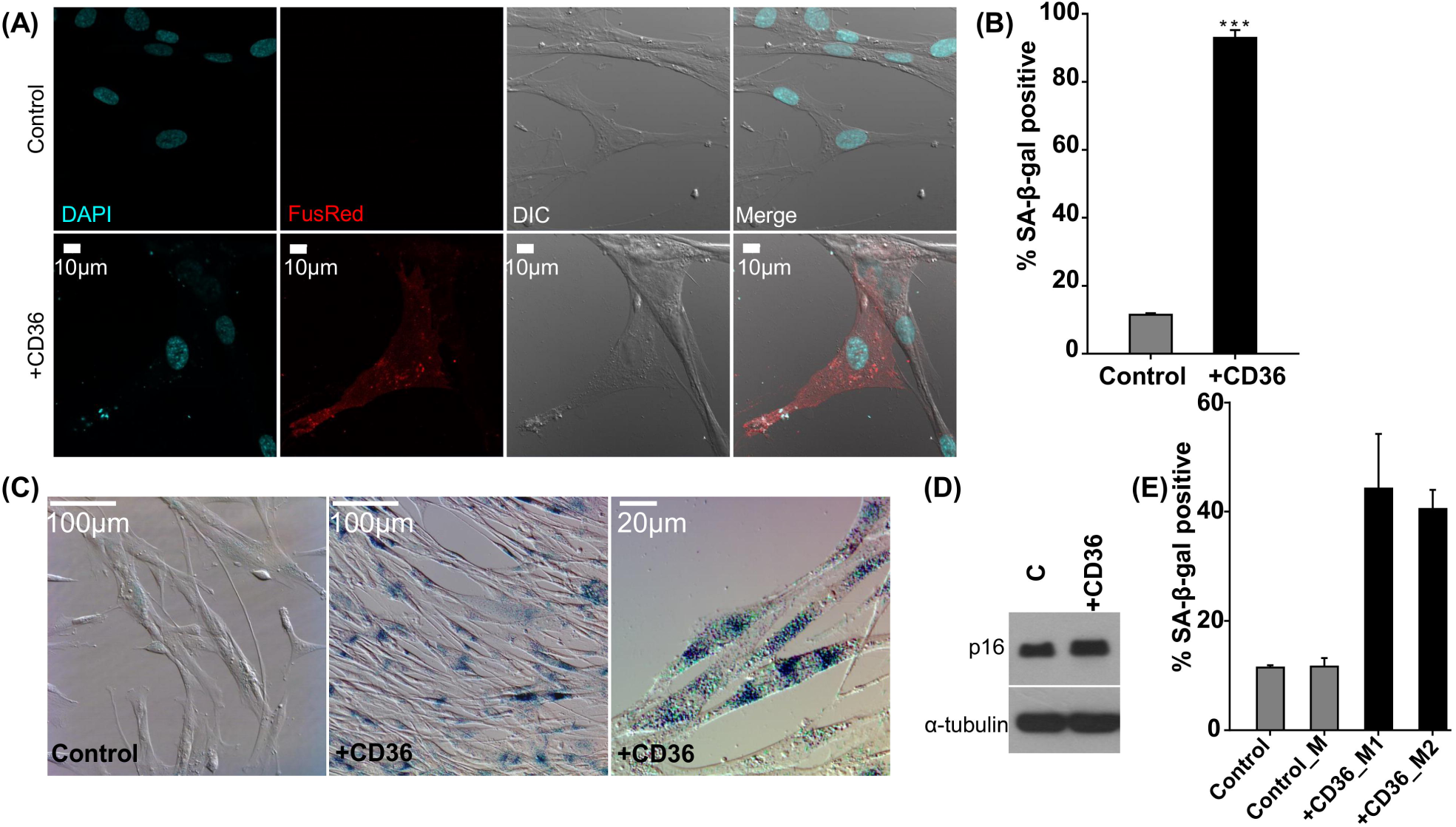
*CD36* overexpression results in senescence-like phenotype. (A) Representative images of control and *CD36*-overexpressing young MRC-5 cells. *CD36 o*verexpression was visualized via the FusionRed tag (scale bar = 10µm). (B-C) Measurement of β-galactosidase activity in control and *CD36* overexpressing MRC-5 cells. A significant increase of SA-β-galactosidase positive cells was observed during *CD36* overexpression for which representative images are shown (scale bar =100µm for control and center CD36 images, and 20µm for furthest to the right CD36 image). Data shown as means ± standard deviation, *** p < 0.001. (D) Representative western blot showing p16 levels in control and *CD36* overexpressing young MRC-5 cells.α-tubulin was used as a loading control. We observe a modest increase in p16 levels. (E) Measurement of β-galactosidase activity in young MRC-5s that were treated with conditioned media. “Control” represents cells in standard growth medium, “Control_M” represents cells in medium from control wells during transfection, “+CD36_M1” represents cells in medium from *CD36* overexpressing cells 48h after transfection, and “+CD36_M2” represents cells in medium from *CD36* overexpressing cells 96h after transfection. An increase in SA-β-galactosidase positive cells was observed.

Based on these observations, we hypothesized that CD36 activity may be linked to the SASP. SASP is one of the hallmarks of cellular senescence and is associated with the release of different inflammatory and growth-promoting factors including IL-6, TNF-alpha and growth factor binding proteins (Tchkonia et al. 2013). Recent studies have suggested that senescence is indeed associated with increased extracellular vesicle release containing different proteins and oligonucleotides (Takasugi 2018). Although the composition of the SASP is not fully known and thought to be dependent on cell type and state, the SASP could contribute to the way senescent cells communicate with their surrounding tissue. At the organismal level, studies have suggested that such intercellular crosstalk could be involved in distinct responses including, but not limited to, wound healing and malignancy (Eden et al. 2007). Based on these known connections between activated secretory phenotype and senescence and the profound increase in senescent cells upon CD36 overexpression, we hypothesized that CD36 activation could be implicated in SASP by inducing changes in the lipid composition such that these changes facilitate membrane remodeling, SASP and vesicle release. Activated SASP then further induces a senescence-like phenotype regardless of the overexpression status of CD36.

To test this hypothesis, we harvested growth media from cells that exhibited a senescence-like phenotype after CD36 overexpression at two different time points. Then, we investigated the ability of their media to induce senescence in proliferating cells. Briefly, we overexpressed CD36 for 72 hours, replenished the media with complete growth media and collected the media after 2 days (shown as “**M1**” in **Figure 5E**). At this time, fresh growth media was added, incubated for 2 additional days and harvested (shown as “**M2**” in **Figure 5E**). We then incubated proliferating cells with **M1** and **M2**, and investigated β-galactosidase activity (**Figure 5E**). This resulted in a significant increase in β-galactosidase activity (**Figure 5E**), strongly supporting that as cells develop a senescence-like phenotype upon CD36 overexpression, they also gain the ability to induce a similar senescence phenotype in neighboring cells. Although future studies are needed to dissect the exact molecular mechanism, we hypothesize that CD36 expression might be linked to senescence-inducing SASP formation.

### Changes in lipid composition in senescent MRC-5 cells reflects membrane remodeling

Our results suggested a link between increased levels of CD36 and membrane-related changes that take place during replicative senescence, including the activation of the SASP. Since lipids are the major components of cellular membranes, we envisioned that the membrane remodeling that facilitates SASP and other extracellular vesicle formation should reflect on the changes in the lipidome of senescent cells. To investigate this, we used a targeted lipidomics approach, as we described previously for BJ cells (Lizardo et al. 2017), and studied the changes in lipid composition as MRC-5 cells senesced.

Specifically, we focused on phosphatidylcholines, which are major building blocks of cellular membranes (van Meer et al. 2008). We compared their levels in dividing (young) and senescent (old) cells and found that certain phosphatidylcholine species accumulated significantly as cells senesced (**Figure 6**). This accumulation can be explained partially with the increased levels of CD36. Indeed, we previously linked CD36 activity as one of the major determinants of the changes in lipid composition during replicative senescence (Lizardo et al. 2017). Majority of phosphatidylcholines that accumulated during senescence contained highly unsaturated long fatty acyl chains. Lipid composition of cellular membrane greatly influence their biophysical properties such as elasticity, curvature, and permeability (Janmey & Kinnunen 2006) and subsequently their involvement in different biological processes (Storck et al. 2018). Long chain fatty acyl chain containing lipids have been implicated in membrane deformation (Atilla-Gokcumen et al. 2014; Szafer-Glusman et al. 2008). Based on these, we envision that changes in phosphatidylcholine composition, in particular accumulation of unsaturated species, could be involved in the release of extracellular vesicles and SASP during replicative senescence impacting the biophysical properties of the membrane either on their own or as their peroxidized forms that occur under oxidative stress during senescence. Although the functional role of the fatty acid composition in SASP factors remain to be elucidated, we propose that the changes in lipid composition observed during senescence might be involved in facilitating vesicle release in this process.

**Figure 6.**
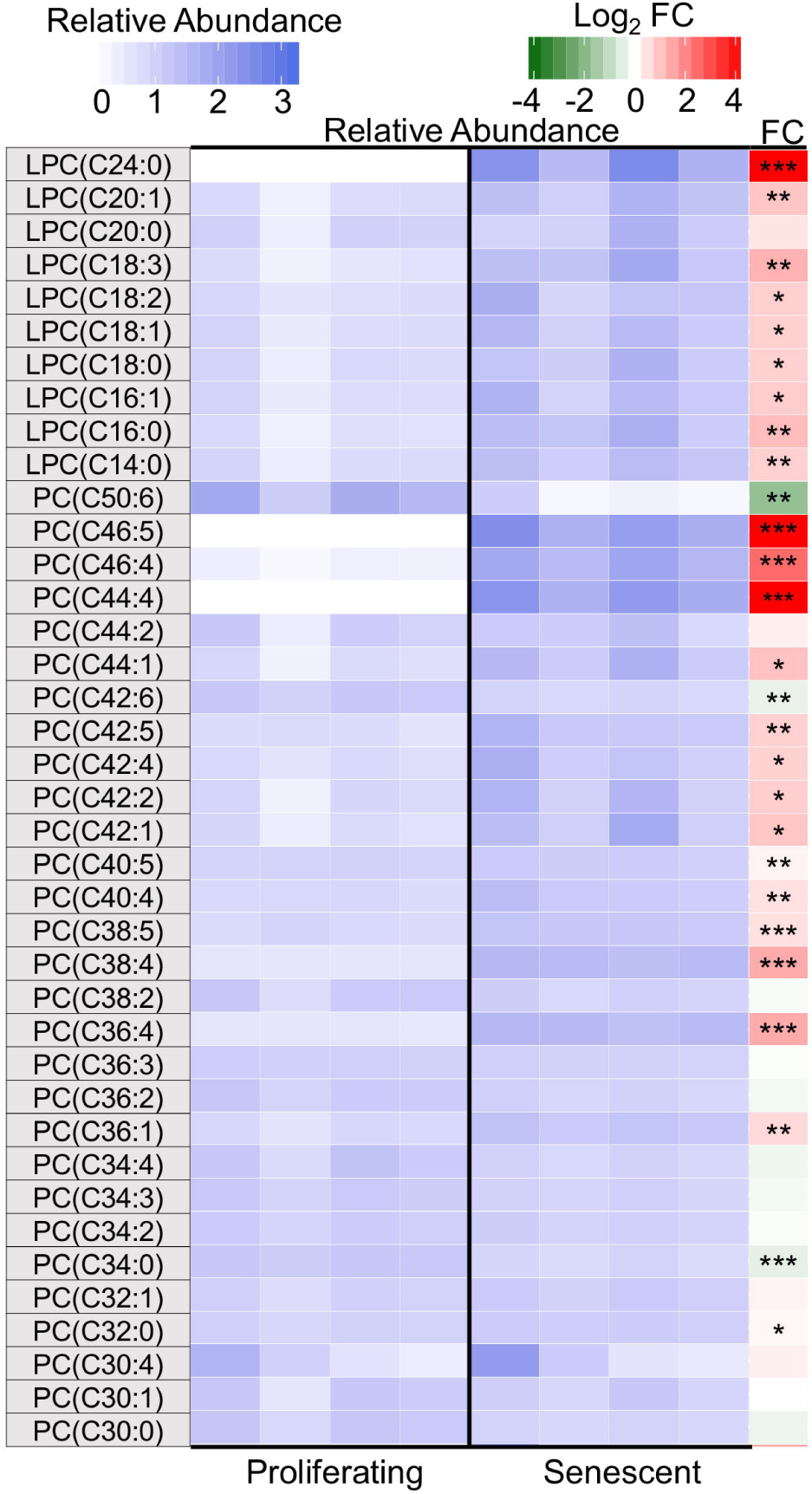
Changes in phosphatidylcholine composition in senescent MRC-5 cells. Targeted analysis of phosphatidylcholines during replicative senescence. Relative abundances were calculated with respect to averaged abundances. Total carbon number on the acyl chains and the degree of unsaturation of the fatty acyl chains in the lipid species are listed, i.e. 38:5 corresponds to 38 carbons and 5 double bonds. Fold change was determined as [Abundance_Senescent_] / [Abundance_Proliferating_] for each lipid species. Abundance is the total ion count for a given ion. Abbreviations: FC: fold change; PC: phosphatidylcholine; LPC, lysophosphatidylcholine. *, p < 0.05; **, p< 0.01; ***, p < 0.001.

## Conclusion

In this study, we report that genes downregulated in senescent cells, which are involved in cell-cycle control, were highly conserved. This finding supports the notion that cellular senescence is tightly regulated. Next, we found that upregulated genes, which are primarily involved in lipid metabolism, were less conserved than downregulated genes. Using these datasets, we conducted an evolutionary transcriptome-wide analysis and identified CD36, as one of the most significantly upregulated lipid-related genes in senescent cells. We showed that overexpression of CD36 in proliferating cells resulted in a senescence-like phenotype. We hypothesized that CD36 overexpression leads to changes in membrane remodeling and plausibly mediate the SASP. Our study employed high-throughput transcriptomics analysis in an evolutionary framework to identify CD36 as a major player in replicative senescence. The exact mechanisms through which CD36 affects cellular senescence, and the evolution of the functional variation in *CD36* will remain a fascinating area of further study.

## Experimental Procedure

### Cells and materials

BJ (human foreskin fibroblasts) and MRC-5 (human lung fibroblasts) cell lines were purchased from the American Type Culture Collection. Minimum Essential Medium (MEM), Eagle’s MEM, penicillin, L-glutamine, and trypsin were acquired from Corning. Fetal Bovine Serum (FBS) was purchased from Sigma-Aldrich. FusionRed-CD36-C10 plasmid (#56104) was obtained from Addgene (deposited by the Davidson lab). Lipofectamine 3000 and ProLong Gold with DAPI were purchased from Invitrogen. Anti-CDKN1A (p21) (# 2946S) and Senescence-associated β-galactosidase staining kit (# 9860) were purchased from Cell Signaling. Anti-CDKN2A (p16) (# ab108349) was purchased from Abcam. Reagents and buffers needed for RNA extraction were obtained from Qiagen. Plasmids were extracted using Omega Bio-Tek E.Z.N.A. FastFilter Plasmid Midi Kit (#D6905-03). LB broth and granulated agar were purchased from BD Biosciences.

### Cell culture and transfection conditions

BJ cells were cultured in MEM while MRC-5s were cultured in Eagle’s MEM supplemented with 1.5 g/L sodium bicarbonate, non-essential amino acids, and sodium pyruvate. Growth medium was also supplemented with 2 mM L-glutamine, 10% (v/v) FBS and 1% (v/v) penicillin-streptomycin. They were grown as monolayer culture at 37 °C and 5% CO_2_ atmosphere under sterile conditions. Cells were routinely checked for mycoplasma contamination. The cumulative population doubling was calculated at each splitting using the formula log2 (D/D_0_), where D is the density of cells during harvesting and D_0_ is the density of cells at seeding. Transfections were performed using Lipofectamine 3000 according to the manufacturer’s instructions. Briefly, 0.30 × 10^6^ cells were seeded in 6-well plates in medium supplemented with 10% (v/v) FBS. After 24 hours, DNA-lipid complex was added to the cells. The transient expression of FusionRed-CD36 was detected using a Leica DMI6000B inverted microscope equipped with a Leica-DFC3000G camera using the LAS AF software at an excitation wavelength of 580 nm for 72 hours after transfection. After transfection, cells were used for imaging or stained for β-galactosidase activity.

### Senescence-associated β-galactosidase (SA-β-gal) activity

Approximately 5.0 × 10^4^ cells were seeded in 24-well plates or 0.30 × 10^6^ cells were plated in 35mm glass bottom dishes. Growth medium was supplemented with 10% (v/v) FBS. After 24 hours of seeding, cells were transfected using Lipofectamine 3000 according to the manufacturer’s instructions. At end of transfection, cells were washed with pre-warmed phosphate buffered saline (PBS). SA-β-gal activity was assessed using the Senescence-associated β-galactosidase staining kit (Cell Signaling) according to manufacturer’s instructions. Briefly, cells were fixed using fixation buffer for 10 minutes. The cells were then incubated overnight at 37 °C in freshly prepared β-galactosidase staining solution which had a final pH of 6.0. At the end of the incubation, cells were washed with PBS and observed under a light microscope in order to determine the number of SA-β-gal positive cells (n=4 for each condition). For control wells, 0.25 × 10^4^ cells were plated 48 hours before staining for β-galactosidase, treated with Lipofectamine reagent for 24 hours and then stained for β-galactosidase. The assay was carried out in three independent experiments where total n ≥ 150 cells for each group. To assess senescence state as fibroblasts stopped dividing, we plated ~4.0 × 10^4^ of old and young cells and carried out the staining as described above.

### Isolation of total RNA and preparation of transcriptomics data

We harvested approximately 2.0 × 10^6^ cells at “young” (BJ= PD15, MRC-5= PD31), “mid-population”(BJ= PD31, MRC-5= PD46), and “old” (BJ= PD44, MRC-5= PD67) stages and extracted total RNA using the RNeasy Mini kit (Qiagen) according to manufacturer’s instructions. An on-column DNase digestion using Qiagen RNase-Free DNase was carried out for RNA purification. RNA quality and yield were assessed by a Nanodrop-1000 spectrophotometer (Thermo). Concentrations ranged from 50-609 ng/μL. Illumina Sequencing (150bp paired-end, ~50M/reads per sample) was performed by the New York Genome Center (http://www.nygenome.org/) using three biological replicates per condition for each cell line. We then analyzed these datasets using Kallisto (Bray et al. 2016) and R package DESeq2 (Love et al. 2014). DESeq2 assumes that all the genes measured in the same experiment have similar variance across replicates and reduces the noise resulting from the low count number of a gene by correcting the variance of each gene. DESeq2 calculates the fold change of each gene using the Wald test and a correction for multiple hypotheses. The entire gene expression of the 18 samples are provided in **Table S4**.

### The filtering pipeline to obtain genes associated with replicative senescence

Of the 57,445 genes we investigated, including protein-coding genes, pseudogenes, antisense RNAs and long non-coding RNAs, we first extracted genes whose expression significantly differed between young and old for BJ and MRC-5 cells. Statistical significance was defined as *p*<0.001, the adjusted value for multi hypotheses tests. The list of core senescence genes are provided in **Table S1**.

### Clustering analysis

We conducted a clustering analysis by calculating Spearman’s correlation of all the reported 57,445 gene expressions between each combination of the 18 samples (BJ_Young, BJ_Mid-population, BJ_Old, MRC-5_Young, MRC-5_Mid-population, MRC-5_Old), without any *a priori* clustering.

### Functional characterization of the genes involved in senescence

We conducted GO (Ashburner et al. 2000; The Gene Ontology Consortium 2017) analysis with Gorilla (Eden et al. 2009; Eden et al. 2007). The target input was the genes upregulated and downregulated in old cells, respectively and the background input was all other protein-coding genes.

### Evolutionary conservation analysis of genes involved in replicative senescence

The pLI (the probability of being loss-of-function intolerant) value was used as a standard for the evolutionary conservation of genes of interest. The pLI value is available at The Exome Aggregation Consortium (ExAC), http://exac.broadinstitute.org/ (Lek et al. 2016). Also, b-value (the index of background selection between primate species) was used as a standard for selection (McVicker et al. 2009).

### Media add-back

Young MRC-5 cells were transfected as described above. After 72 hours of transfection, media containing DNA-lipid complex was removed, cells were washed with PBS and supplemented with complete growth medium. The cells were then incubated at 37 °C and 5% CO_2_ atmosphere under sterile conditions for 48 hours. At the end of incubation, conditioned medium from control and CD36 transfected cells was collected and stored at 4 °C. The procedure was repeated a second time to collect conditioned medium 96 hours after transfection. Approximately 5.0 × 10^4^ young MRC-5 cells were seeded in 24-well plates. 24 hours after seeding, a PBS wash was carried out and conditioned medium was added to the corresponding wells. Cells were maintained in conditioned medium for 48 hours and assessed for β-galactosidase activity as described above.

### Image acquisition and processing

Cells (~5.0 × 10^4^/coverslip) were plated on sterile coverslips in 24-well plates or in 35mm glass bottom dishes (0.30 × 10^6^/dish) for live cell imaging or to stain for β-galactosidase. After 24 hours of seeding cells were transfected using Lipofectamine 3000 according to the manufacturer’s instructions. At end of transfection, cells grown on coverslips were washed with pre-warmed PBS and fixed in 3.7% formaldehyde for 20 minutes. Coverslips were then gently rinsed with PBS (x3) and subsequently mounted using ProLong Gold with DAPI antifade reagent. Live cell and β-galactosidase images were acquired on a LSM710 confocal microscope (Carl Zeiss, Oberkochen, Germany). Image analysis was performed using the Fiji/ImageJ software version 2.0.0-rc-49/1.51a. Images are representative of images taken from at least two independent experiments.

### Western blot analysis

About 0.50 × 10^6^ cells were pelleted and lysed with mammalian protein extraction reagent containing a protease inhibitor cocktail. Cellular debris was removed by centrifugation and the supernatant was used to measure protein concentration by Bradford Assay. Equal amounts of proteins were resolved by sodium dodecyl sulfate-polyacrylamide-gel electrophoresis and then transferred onto polyvinyl difluoride membranes using a wet transfer cell. Membranes were blocked in 5% non-fat dry milk in TBS-Tween (0.2% Tween in Tris-buffered saline) for 1 hour. Membranes were then washed four times (8 minutes each) in TBS-tween followed by incubation overnight at 4 °C with primary antibody. Antibodies were diluted in AbDil-tween according to manufacturer’s instructions. Once the incubation with primary antibody was complete, membranes were washed four times (8 minutes each) in TBS-tween followed by incubation with horseradish peroxidase-conjugated anti-mouse or anti-rabbit secondary antibody diluted in 5% non-fat dry milk in TBS-tween for 2 hours at room temperature. Once the incubation with secondary antibody was complete, membranes were washed in TBS-tween four times (8 minutes each) and developed using the Super Signal West Pico kit (ThermoFisher Scientific) according to manufacturer’s guidelines. α-tubulin was used as a loading control.

### Plasmid extraction

CD36 plasmid was received as a bacterial stab (addgene) and grown in 10mm × 20mm petri dishes overnight at 37 °C using the appropriate antibiotics. Single colonies were then picked and grown overnight in 50 mL culture using standard procedures. Cells were then pelleted by ultracentrifugation at 4 °C. Plasmid DNA was isolated using the E.Z.N.A.^®^ FastFilter Plasmid Midi Kit (OMEGA Bio-Tek) following the manufacturer’s instructions. Plasmid DNA concentration and integrity was measured using Thermo Nanodrop-1000 spectrophotometer. Plasmid DNA was then aliquoted into sterile microcentrifuge tubes and stored at -80 °C.

### Lipid extraction and analysis

Preparation of lipid extracts, sample normalization, and liquid chromatography mass spectrometry data acquisition and analysis were performed as previously described (Lizardo et al. 2017).

## Acknowledgements

This study was supported by the Astellas Foundation for Research on Metabolic Disorders (to M.S.). We would like to thank Alan Siegel at the UB North Campus Imaging Facility for help with image acquisition (National Science Foundation Major Research Instrumentation Grant # DBI 0923133). We would like to thank Dr. Mehmet Somel for his comments on the manuscript.

## Author Contributions

All authors participated in the preparation of the manuscript. OG and GEAG performed study conception and design. DL and AM carried out lipidomics analysis and characterization of CD36 overexpressing cells. MS, RT and OG analysed the transcriptomics data.

## Supplementary FIgures

**Figure S1**. Characterization of replicative senescence in BJs and MRC-5s. (A). BJs and MRC-5s were grown under standard culture conditions until the naturally reached their proliferative capacity. Yellow points: MRC-5s, blue points: BJs. For transcriptomics analysis pellets were collected at PD31 as “young”, PD46 as “mid-population” and PD67 as “old” MRC-5s. For BJs, PD15 were collected as “young”, PD31 as “mid-population” and PD44 as “old”. y-axis represents the cumulative population doubling and x-axis represents the days cells were maintained in culture. (B) Measurement of βeta-galactosidase activity in BJ and MRC-5 cells during replicative senescence. A significant increase of SA-β-galactosidase positive cells was observed as the cells senesced. Data shown as means ± standard deviation, *** p < 0.001.

**Figure S2**. Live MRC-5 cell tile scan. Representative images of control and *CD36* transfected live cells young MRC-5 cells. As illustrated by the transient expression of FusionRed-CD36 (excitation wavelength = 580 nm), the transfection efficiency was ~10% of young MRC-5 cells. The tile scan was collected by adjoining single images. Scale bar = 100µm.

**Table S1**. The master table for evolutionary transcriptomics analyses for all protein coding genes analyzed in this study. GeneID and Gene columns indicate the ENSEMBL and common names for each gene, respectively. pLI, pRec, pNull values are measures of tolerance to loss of function, retrieved from The Exome Aggregation Consortium (ExAC), http://exac.broadinstitute.org/ (Lek et al. 2016). The next four columns indicate the log_2_ fold change and multiple hypotheses corrected *p-*value for the expression change from young to old in BJ and MRC-5 cells. B.value column shows the index of background selection between primate species. These values were retrieved from (McVicker et al. 2009). The GWAS column indicates whether or not these genes were associated in aging-related phenotypes (https://www.ebi.ac.uk/gwas/).

**Table S2**. Genes that showed significant (adjusted *p*<0.0001) fold change of expression between young and old cells in both BJ and MRC-5 cell lines are provided. GeneID and Gene columns indicate the ENSEMBL ID and common names for each gene, respectively. Column Annotation shows the gene annotations. Next two columns show the mean expression in BJ and MRC-5 cells. The Direction column indicates whether the gene is downregulated or upregulated during replicative senescence.

**Table S3**. The enriched GO categories of genes upregulated and downregulated in old cells. Downregulated genes (columns A-O) and upregulated genes (columns T-AH) are listed.

**Table S4**. The gene expression (relative read depth) of a total of 18 samples (triplicates for three time points for BJ and MRC-5 cell lines). BY: young BJs, BM: mid-population BJs, BO: old BJs, MY: young MRC-5s, MM: mid-population MRC-5s, MO: old MRC-5s.

